# Triple layered QSAR Studies on Substituted 1,2,4-Trioxanes as potential antimalarial agents: Superiority of the Quantitative Pharmacophore-Based Alignment, Over Common Substructure based Alignment

**DOI:** 10.1101/468959

**Authors:** Amit K. Gupta, Anil K. Saxena

**Affiliations:** Medicinal and Process Chemistry Division, C.S.I.R.-Central Drug Research Institute, Lucknow, 226001, India

**Keywords:** CoMFA, CoMSIA, HQSAR, substituted 1,2,4-trioxanes, Pharmacophore, Drug design

## Abstract

The present study reports the utilization of three approaches viz Pharmacophore, CoMFA, CoMSIA and HQSAR studies to identify the essential structural requirements in 3D chemical space for the modulation of the antimalarial activity of substituted 1,2,4 trioxanes. The superiority of Quantitative pharmacophore based alignment (QuantitativePBA) over global minima energy conformer-based alignment (GMCBA) has been reported in CoMFA and CoMSIA studies. The developed models showed good statistical significance in internal validation (q^2^, group cross-validation and bootstrapping) and performed very well in predicting antimalarial activity of test set compounds. Structural features in terms of their steric, electrostatic, and hydrophobic interactions in 3D space have been found important for the antimalarial activity of substituted 1,2,4-trioxanes. Further, the HQSAR studies based on the same training and test set acted as an additional tool to find the sub-structural fingerprints of substituted 1,2,4 trioxanes for their antimalarial activity. Together, these studies may facilitate the design and discovery of new substituted 1,2,4-trioxane with potent antimalarial activity.

## 1. Introduction

In spite of worldwide efforts to combat malaria, it still kills approximately one million people, mostly children, each year [1,2]. There is no fully effective prophylactic vaccine against malaria till date [3,4] and the major problem in the chemotherapy of malaria is the development of resistance of the *Plasmodium falciparum* parasites to many of the standard quinoline antimalarial drugs like chloroquine [5]. The discovery of artemisinin, extracted from the plant *Artemisia annua*, has opened a new era in the malarial chemotherapy. Artemisinin and its more potent analogues *viz.* artemether, arteether and artesunic acid represent the endoperoxide class of compounds which are highly active against both chloroquine-sensitive and chloroquine-resistant strains of *P. falciparum* [6]. The WHO-recommended artemisinin combination therapy (ACT) is the best option available till date for the chemotherapy of malaria [7]. The ligand based approaches like three-dimensional Quantitative Structure Activity Relationship (3D-QSAR) studies have been quite useful in identifying the essential structural requirements for biological activity of the compounds where the 3D structure of the exact target is unknown[8, 9]. Therefore, considering the importance of artemisinin and its analogues as potent class of antimalarial drugs effective against the multidrug-resistant *P. falciparum* strains and unavailability of the exact target for this class of molecule [10] we have earlier reported the Discovery studio (DS)[11] based quantitative pharmacophore model utilizing this class of molecules[12]. The DS based pharmacophore models are more computationally intensive, as they consider many conformations (number ≤ 255) of each molecule to generate the QSAR equations [13], but they give the minimum essential structural requirements for the activity in terms of favorable regions and do not give any information about the features that diminishes the biological activity. The successful application of CoMFA, CoMSIA technique to understand the effect of contrast of structural requirements in 3D chemical space has been reported by many research groups in recent past [14,15]. Thus, on our next move we focused on to the less computationally intensive CoMFA, CoMSIA models on the same dataset which not only provide the information about the favorable regions but also give the information about the unfavorable regions in defining the potency. Structural alignment is perhaps the most subjective, yet critical, step in CoMFA study. In our earlier studies the global minima energy conformer-based alignment (GMCBA) had shown better results than docked conformer-based alignment (DCBA), and co-crystallized conformer-based alignment (CCBA) [16]. Further, we had also reported the superiority of qualitative pharmacophore based alignment (QualitativePBA) over GMCBA where the co-crystallize structure of the molecule is unknown [17]. Now, we herein report the superiority of quantitative pharmacophore based alignment (QuantitativePBA) over GMCBA in terms of statistical significance. Besides the knowledge gained from the CoMFA, CoMSIA studies in terms of favorable and unfavorable features in 3D space to regulate the antimalarial activity of this class of compounds, the HQSAR studies based on the same molecular conformations of the training and test sets were also performed to generate the molecular fingerprints for the structures of the artemisinin derivatives relevant to their antimalarial activity. The HQSAR offers the ability of rapid and easy generation of high statistical quality QSAR models [18]. The premise of Hologram QSAR (HQSAR) is based on the assumption that the structure of a molecule is a key determinant (fingerprint) of the biological activity. The HQSAR studies however use an extended form of fingerprint, known as a "molecular hologram", which encodes more information in terms of branched and cyclic fragments including their stereochemistry, than the traditional 2D fingerprint. Together, the resulting three layered QSAR models will help to better understand the role of different chemical features in governing antimalarial activity of substituted 1,2,4-trioxanes and may serve as a tool for developing more potent antimalarial agents.

## 2. Method and Material

### 2.1. Biological activity

The QSAR studies were performed using eight series of substituted 1, 2, 4-trioxanes comprising 88 artemisinin analogues (activity ranges from 1.4nM to 2000nM) reported in literature [19-24]. The homogeneity of the biological assays is one of the important aspects in QSAR study therefore, the dataset was collected from the same research group following the same biological testing protocol. It has been suggested that the generated models should be tested on a sufficiently large test set to establish a reliable QSAR model [25] therefore, the molecules were rationally divided into training set of 45 and the test set of 43 compounds in such a way that both sets cover the structural diversity of following eight different chemical prototypes and entire range of biological activity (Figure 1 and Table 1).

**Table 1.**
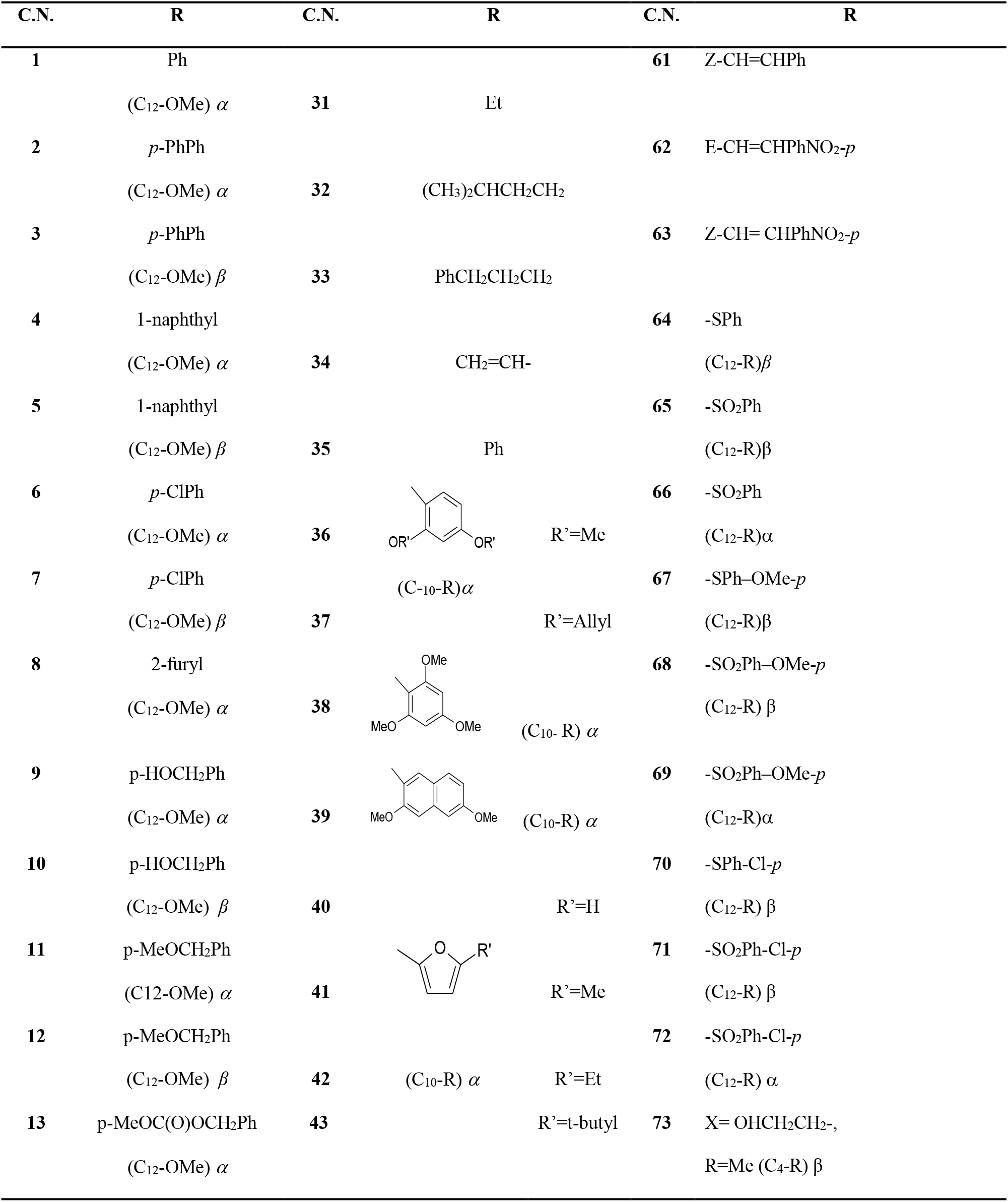
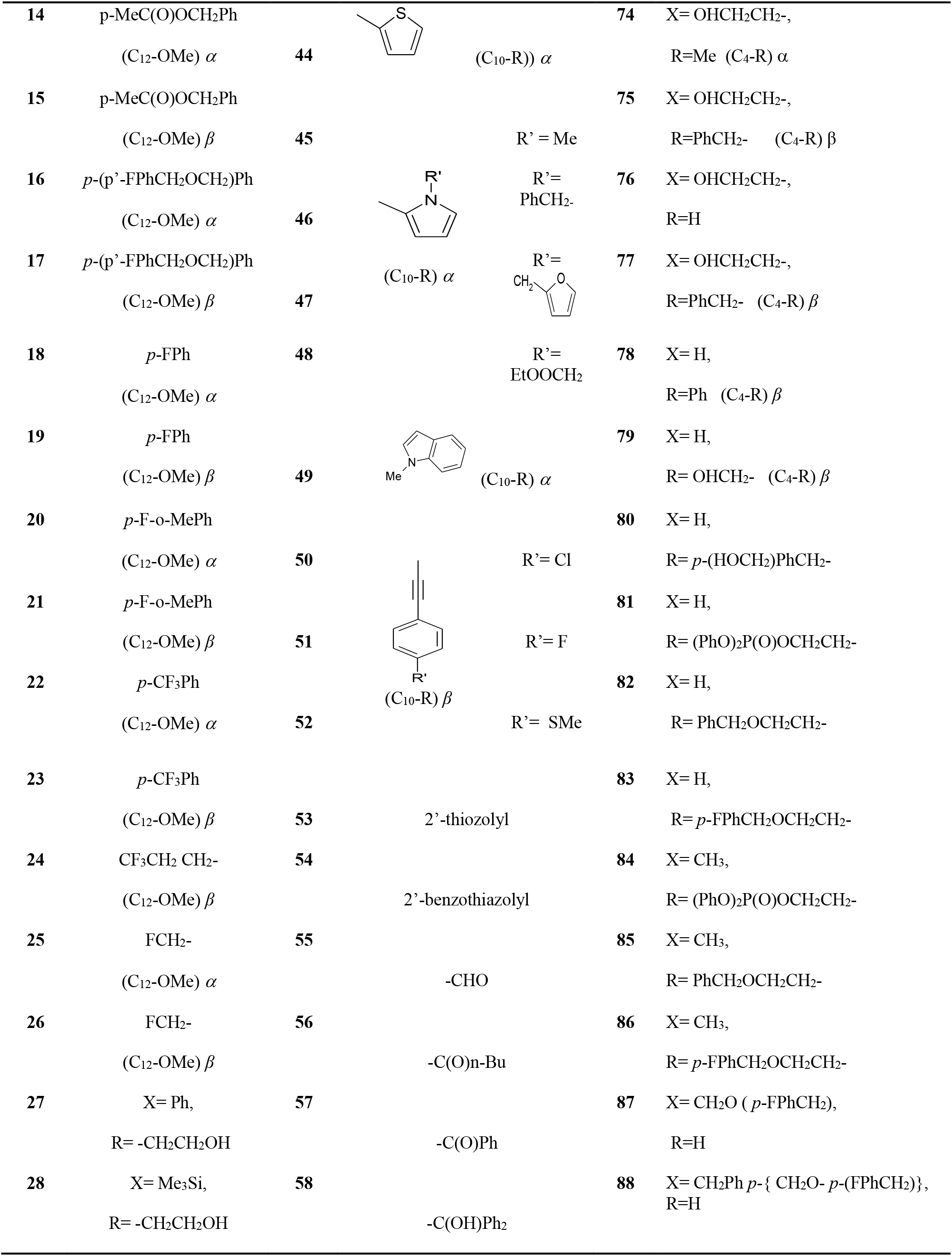
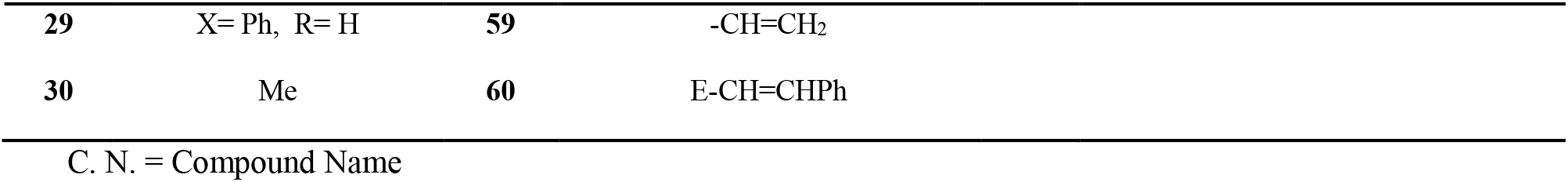
Structures of the molecules used in training and test set

**Figure 1.**
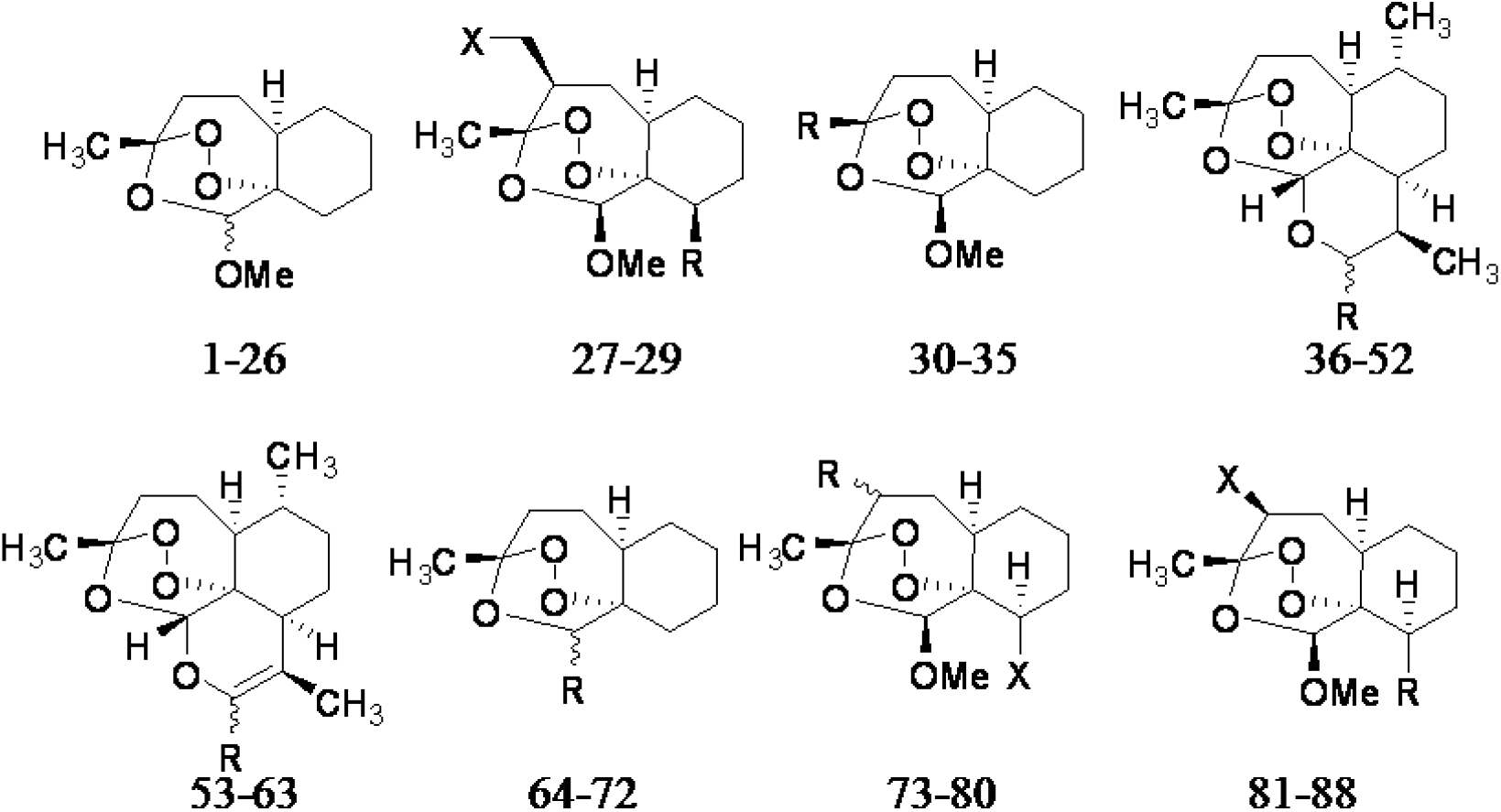
Structures of compounds used in training and test set.

### 2.2 CoMFA and CoMSIA studies

#### 2.2.1 Computational approach and Molecular alignment

The CoMFA and CoMSIA molecular modeling studies were performed using SYBYL software [26] running on a Silicon Graphics Octane R12000 workstation. In ‘GMCBA’ method, the 3D structures of the molecules to be analyzed are aligned to a suitable conformational template, which is assumed to adopt a ‘‘bioactive conformation’’. Hence, in this case the molecular structures of all the compounds were drawn using the most active compound **40** as a template in SYBYL6.9 where the partial charges were calculated using Gasteiger-Hückel method and geometry optimized using Tripos force field with a distance-dependent dielectric function and energy convergence criterion of 0.001 kcal/mol Å using 1000 iterations and standard SYBYL settings. The conformational search was performed using multi-search method with the following settings: maximum cycles (400), maximum conformers (400), energy cutoff (70 kcal/mol), maximum rms gradient (3.0) tolerance (0.40), and number of hit (12). The minimum energy conformations thus obtained were used in the GMCBA analysis. The substructure (shown in blue color) of the most active compound **40** (Figure 2A) was used as a template for molecular alignment. Whereas, in the QuantitativePBA analysis the earlier reported quantitative pharmacophore based alignment of all the 88 artemisinin derivatives was exported to SYBYL6.9 interface for CoMFA and CoMSIA studies (Figure 2C). The partial charges for all the compounds were calculated using the Gasteiger-Hückel method. The overall alignment of the training set molecules for the GMCBA and QuantitativePBA method has been shown in Figure 2B and 2D.

**Figure 2.**
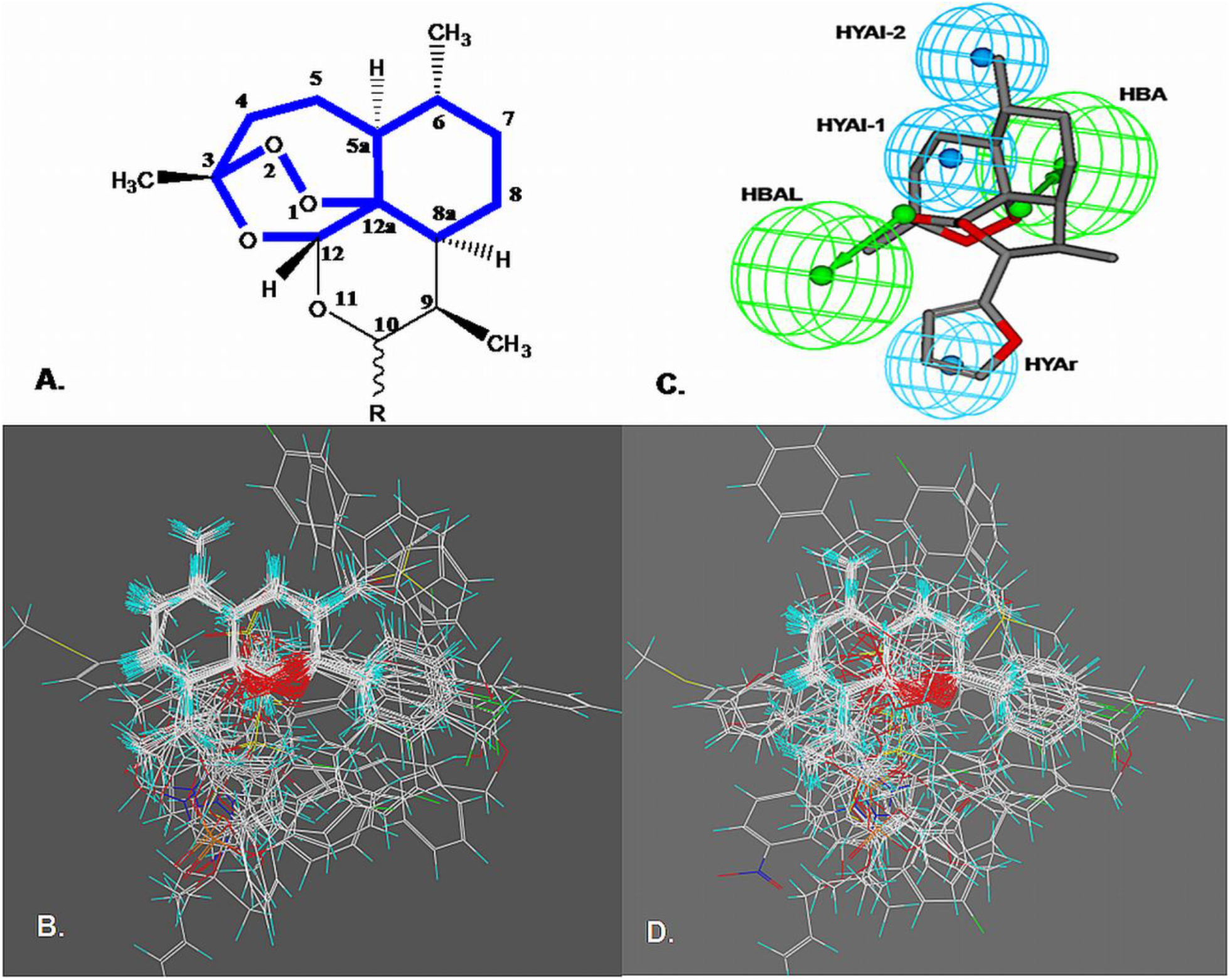
The overall alignment of the training set molecules used in the 3D-QSAR study, (**A**) global minima energy conformer-based alignment, (**B**) Quantitative pharmacophore based alignment.

#### 2.2.2 CoMFA

The CoMFA methodology was first reported by Cramer *et al*.[27] through which a three-dimensional QSAR model can be derived for a set of ligands by sampling the steric and electrostatic fields around them with respect of their biological activity. In the present study the aligned molecules of the training set were positioned in a 3D cubic lattice with a grid spacing of 2.0 Å in x, y and z directions for deriving the CoMFA fields. The steric (Lennard–Jones potential) and electrostatic (Columbic with 1/r dielectric) fields were calculated at each lattice point using Tripos force field and a distance dependent dielectric constant of 1.0. An sp^3^-hybridized carbon with a +1.0 charge and a radius of 1.52 Å was used as a probe to calculate various steric and electrostatic fields. An energy cutoff value of 30 kcal/mol was applied to avoid too high and unrealistic energy values inside the molecule.

#### 2.2.3 CoMSIA

The CoMSIA technique is based on the molecular similarity indices with the same lattice box used for the CoMFA calculations [28]. It is considered superior to CoMFA technique in certain aspects such as the results remain unaffected to both, region shifts as well as small shifts within the alignments, it does not require steric cutoffs and more intuitively interpretable contour maps. In the present study, five different similarity fields *viz.* steric, electrostatic, hydrophobic, H-donor and H-acceptor were calculated using the standard settings of CoMSIA (Probe with charge +1, radius 1 Å and hydrophobicity +1, hydrogen-bond donating +1, hydrogen-bond accepting +1, attenuation factor of 0.3 and grid spacing 2 Å).

#### 2.2.4 Partial least squares (PLS) and Predictive r_2_ analysis

PLS is used to correlate sirt1 activity with the CoMFA and CoMSIA values containing magnitude of steric, electrostatic and hydrophobic potentials. Leave one out (LOO) validation was utilized as tool for determining the predictability of the developed model. The full PLS analysis was carried out with a column filtering of 2.0 kcal/mol to speed up the calculation and reduce the noise. The predictive r^2^ value is based on the only test set molecules which may be define as

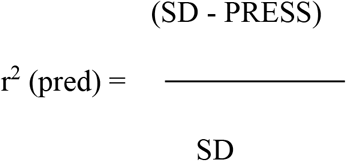

where SD is the sum of squared deviation between the biological activities of the test set molecule to the mean activity of the training set molecules while PRESS is the sum of squared deviations between the observed and the predicted activities of the test molecules.^36^

### 2.3 Hologram QSAR (HQSAR) studies

A molecular hologram development for a number of structures can yield a data matrix of dimension N x L. N is the number of compounds analyzed and L is the length of the molecular hologram. The PLS method is used to build a statistical model which relates the molecular hologram descriptors to an experimental property such as pIC50. The combinations of different fragments viz. atomic numbers (A) to distinguish the atom types, bond types (B) to distinguish the bond types, atomic connections (C) to consider the hybridization states of the atoms in a fragment, hydrogen (H) for inclusion of H-atoms, donor and acceptor (DA) for donor and acceptor atoms, and chirality (Ch) were accessed for all the 88 artemisinin derivatives during the hologram generation. This choice was based on the structural features likely to be important in the data set being analyzed. The non-cross-validated regression coefficient (r^2^), cross-validated regression coefficient value (q^2^) and standard error of prediction (SEE) for different combinations of fragments were analyzed to select the best combinations as a molecular hologram and the HQSAR model.

## 3. Results and discussion

### 3.1 CoMFA and CoMSIA studies

Since, the CoMFA and CoMSIA are highly sensitive to the relative alignment of molecules, it was important to determine the best alignment rule for these molecules having varying structures. Among the QuantitativePBA and GMCBA methods for the alignment for all the 88 artemisinin derivatives, the best alignment was selected on the basis of statistical parameters obtained both for CoMFA and CoMSIA models listed in Table 2. Since the QuantitativePBA based alignment method gave the model with best statistics and predictive values, this alignment was further used for systematic CoMFA, and CoMSIA studies.

**Table. 2.** PLS statistics of CoMFA (TS) and CoMSIA (SEHA) models.

| Parameters |  | GMCBA |  | QuantitativePBA |  |
| --- | --- | --- | --- | --- | --- |
|  |  | CoMFA (TS) | CoMSIA (SEH) | CoMFA (TS) | CoMSIA (SEH) |
| $q^2$ | | 0.341 | 0.331 | 0.721 | 0.725 |
| $S_{press}$ | | 0.589 | 0.593 | 0.383 | 0.410 |
| $r^2$ | | 0.527 | 0.499 | 0.834 | 0.875 |
| SEE |  | 0.499 | 0.513 | 0.292 | 0.257 |
| F |  | 47.857 | 42.902 | 222.982 | 150.728 |
| N |  | 1 | 1 | 1 | 1 |
| Fractions | S | 0.534 | 0.228 | 0.352 | 0.152 |
|  | E | 0.466 | 0.413 | 0.648 | 0.455 |
|  | H |  | 0.359 |  | 0.393 |
| $r^2_{bs}$ (20 runs) | | 0.624 | 0.570 | 0.821 | 0.879 |
| $SD_{bs}$ | | 0.432 | 0.469 | 0.303 | 0.239 |
| $r^2_{CV(mean)}$ (20 groups) | | 0.341 | 0.333 | 0.724 | 0.741 |
| $r^2_{pred}$ | | 0.482 | 0.482 | 0.635 | 0.713 |

$q^2$ = leave one out cross-validation correlation coefficient; $r^2$ = conventional correlation; SEE = standard error of estimate; F = degree of freedom; N = optimal number of component; $r^2_{bs}$ = bootstrapping correlation; $SD_{bs}$ = bootstrapping standard deviation; $r^2_{CV(mean)}$ = group cross-validation; TS = Tripos standard; SEH = Steric, electrostatic, and Hydrophobic.

#### 3.1.1 CoMFA analysis

The QuantitativePBA method based CoMFA analysis revealed a cross validated q^2^ of 0.721 for 1 principal components and a non cross validated conventional r^2^ of 0.834, F value of 222.982 and standard error of estimate SEE of 0.292. To further assess the robustness of the models, bootstrapping analysis (20 runs) was performed and the observed r^2^_bs_ of 0.821 (SD_bs_= 0.036), further strengthened the model. In addition to LOO, a group cross-validation for 20 runs was also carried out to assess the internal predictive ability of the model and the observed mean r^2^_CV_ of 0.724 (TS) was indicative of high internal predictivity of the model (Table 2).

#### 3.1.2 CoMSIA analysis

The CoMSIA model having steric(S), electrostatic (E), and hydrophobic (H) descriptors gave the highest q^2^ value of 0.725 for 1 component with a conventional non cross-validated r^2^ value 0.875, F value 150.728 and standard error of estimate SEE 0.257. To further assess the statistical ability and the robustness of the model, bootstrapping analysis (20 runs) was performed where the observed r^2^_bs_ value 0.879 with very low standard deviation 0.036 indicated the high robustness of the model. Similar to CoMFA, it also showed high internal predictive ability (mean r^2^_cv_ value 0.741) for 20 runs.

#### 3.1.3 Test Set Validation

The test set validation is the rigorous validation for the model where the activity predictions for compounds not included in the training set are made. The external predictive ability of the generated CoMFA model was evaluated for the test set of 43 molecules where the obtained predictive r^2^ value (r^2^_pred_) of 0.635 further supported the high predictive ability of the generated model (Figure 3A). The predictive pIC_50_ values of the training as well as test set molecules based on the CoMFA model are listed in the Table 3. Similar to the CoMFA model, the CoMSIA models also showed the high external predictive ability (r^2^_pred_) of 0.713 for the external test set (Figure 3B). The observed and predicted activities of the training and test set by the best CoMSIA (SEH) model are shown in Table 3. These results for the test-set compounds provide strong evidence that the CoMFA and CoMSIA models so derived are able to predict well the antimalarial activities of structurally diverse artemisinin derivatives.

**Table 3.**
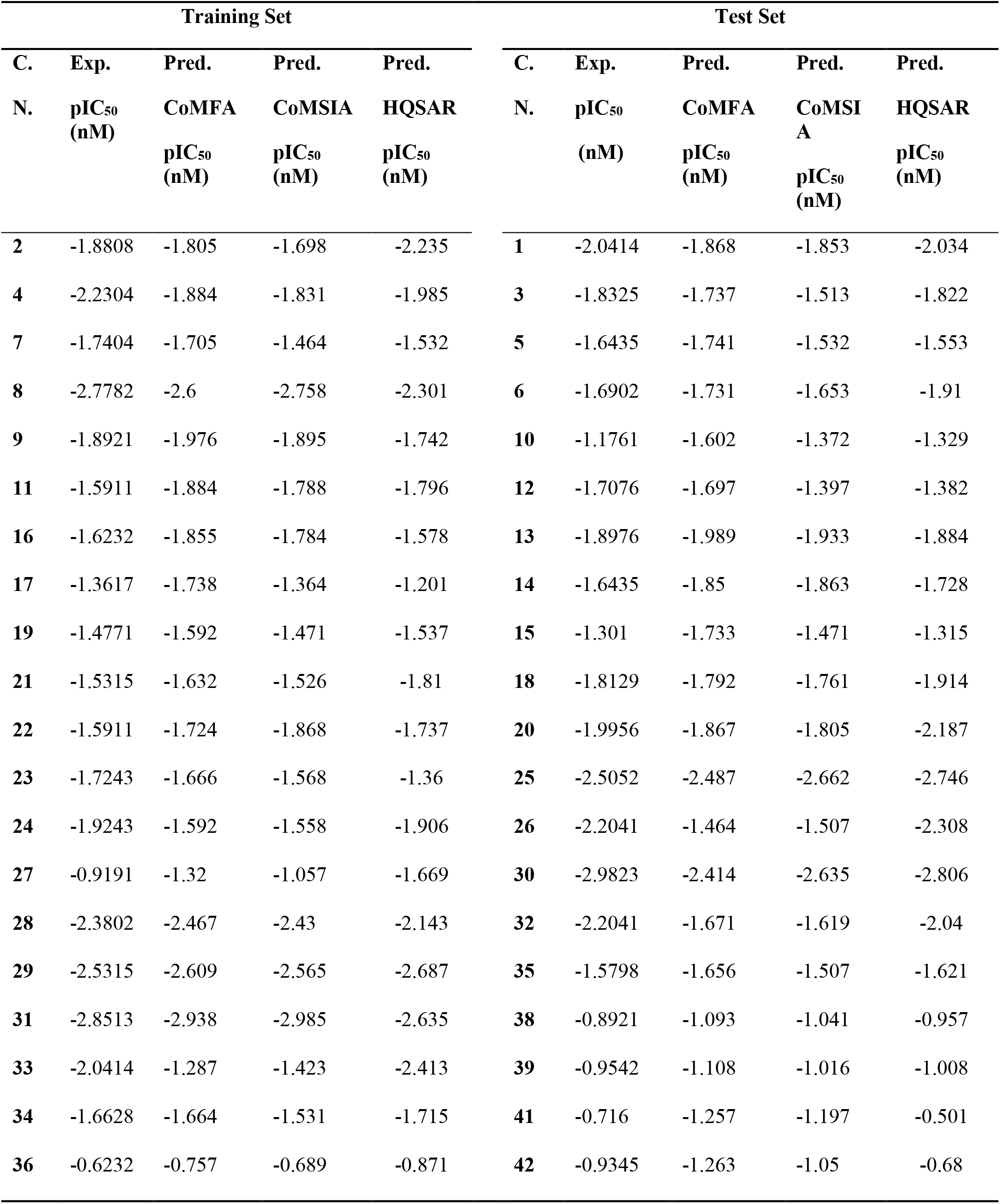
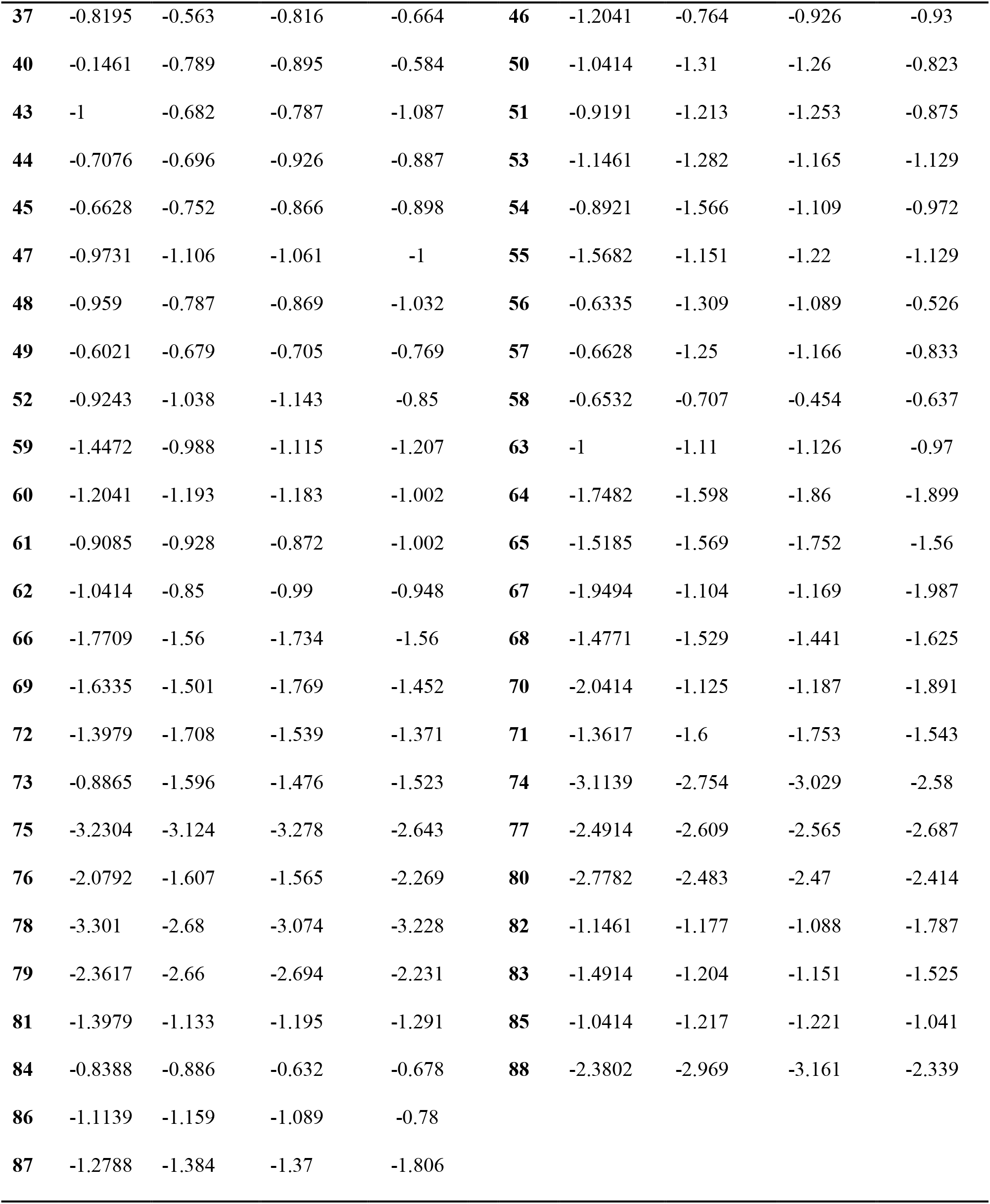
Training set and test set compounds along with their experimental and predicted biological activity.

| Training Set |  |  |  |  | Test Set |  |  |  |  |
| --- | --- | --- | --- | --- | --- | --- | --- | --- | --- |
| C. | Exp. | Pred. | Pred. | Pred. | C. | Exp. | Pred. | Pred. | Pred. |
| N. | pIC<sub>50</sub><br>(nM) | CoMFA<br>pIC<sub>50</sub><br>(nM) | CoMSIA<br>pIC<sub>50</sub><br>(nM) | HQSAR<br>pIC<sub>50</sub><br>(nM) | N. | pIC<sub>50</sub><br>(nM) | CoMFA<br>pIC<sub>50</sub><br>(nM) | CoMSI<br>A<br>pIC<sub>50</sub><br>(nM) | HQSAR<br>pIC<sub>50</sub><br>(nM) |
| **2** | -1.8808 | -1.805 | -1.698 | -2.235 | **1** | -2.0414 | -1.868 | -1.853 | -2.034 |
| **4** | -2.2304 | -1.884 | -1.831 | -1.985 | **3** | -1.8325 | -1.737 | -1.513 | -1.822 |
| **7** | -1.7404 | -1.705 | -1.464 | -1.532 | **5** | -1.6435 | -1.741 | -1.532 | -1.553 |
| **8** | -2.7782 | -2.6 | -2.758 | -2.301 | **6** | -1.6902 | -1.731 | -1.653 | -1.91 |
| **9** | -1.8921 | -1.976 | -1.895 | -1.742 | **10** | -1.1761 | -1.602 | -1.372 | -1.329 |
| **11** | -1.5911 | -1.884 | -1.788 | -1.796 | **12** | -1.7076 | -1.697 | -1.397 | -1.382 |
| **16** | -1.6232 | -1.855 | -1.784 | -1.578 | **13** | -1.8976 | -1.989 | -1.933 | -1.884 |
| **17** | -1.3617 | -1.738 | -1.364 | -1.201 | **14** | -1.6435 | -1.85 | -1.863 | -1.728 |
| **19** | -1.4771 | -1.592 | -1.471 | -1.537 | **15** | -1.301 | -1.733 | -1.471 | -1.315 |
| **21** | -1.5315 | -1.632 | -1.526 | -1.81 | **18** | -1.8129 | -1.792 | -1.761 | -1.914 |
| **22** | -1.5911 | -1.724 | -1.868 | -1.737 | **20** | -1.9956 | -1.867 | -1.805 | -2.187 |
| **23** | -1.7243 | -1.666 | -1.568 | -1.36 | **25** | -2.5052 | -2.487 | -2.662 | -2.746 |
| **24** | -1.9243 | -1.592 | -1.558 | -1.906 | **26** | -2.2041 | -1.464 | -1.507 | -2.308 |
| **27** | -0.9191 | -1.32 | -1.057 | -1.669 | **30** | -2.9823 | -2.414 | -2.635 | -2.806 |
| **28** | -2.3802 | -2.467 | -2.43 | -2.143 | **32** | -2.2041 | -1.671 | -1.619 | -2.04 |
| **29** | -2.5315 | -2.609 | -2.565 | -2.687 | **35** | -1.5798 | -1.656 | -1.507 | -1.621 |
| **31** | -2.8513 | -2.938 | -2.985 | -2.635 | **38** | -0.8921 | -1.093 | -1.041 | -0.957 |
| **33** | -2.0414 | -1.287 | -1.423 | -2.413 | **39** | -0.9542 | -1.108 | -1.016 | -1.008 |
| **34** | -1.6628 | -1.664 | -1.531 | -1.715 | **41** | -0.716 | -1.257 | -1.197 | -0.501 |
| **36** | -0.6232 | -0.757 | -0.689 | -0.871 | **42** | -0.9345 | -1.263 | -1.05 | -0.68 |
| **37** | -0.8195 | -0.563 | -0.816 | -0.664 | **46** | -1.2041 | -0.764 | -0.926 | -0.93 |
| **40** | -0.1461 | -0.789 | -0.895 | -0.584 | **50** | -1.0414 | -1.31 | -1.26 | -0.823 |
| **43** | -1 | -0.682 | -0.787 | -1.087 | **51** | -0.9191 | -1.213 | -1.253 | -0.875 |
| **44** | -0.7076 | -0.696 | -0.926 | -0.887 | **53** | -1.1461 | -1.282 | -1.165 | -1.129 |
| **45** | -0.6628 | -0.752 | -0.866 | -0.898 | **54** | -0.8921 | -1.566 | -1.109 | -0.972 |
| **47** | -0.9731 | -1.106 | -1.061 | -1 | **55** | -1.5682 | -1.151 | -1.22 | -1.129 |
| **48** | -0.959 | -0.787 | -0.869 | -1.032 | **56** | -0.6335 | -1.309 | -1.089 | -0.526 |
| **49** | -0.6021 | -0.679 | -0.705 | -0.769 | **57** | -0.6628 | -1.25 | -1.166 | -0.833 |
| **52** | -0.9243 | -1.038 | -1.143 | -0.85 | **58** | -0.6532 | -0.707 | -0.454 | -0.637 |
| **59** | -1.4472 | -0.988 | -1.115 | -1.207 | **63** | -1 | -1.11 | -1.126 | -0.97 |
| **60** | -1.2041 | -1.193 | -1.183 | -1.002 | **64** | -1.7482 | -1.598 | -1.86 | -1.899 |
| **61** | -0.9085 | -0.928 | -0.872 | -1.002 | **65** | -1.5185 | -1.569 | -1.752 | -1.56 |
| **62** | -1.0414 | -0.85 | -0.99 | -0.948 | **67** | -1.9494 | -1.104 | -1.169 | -1.987 |
| **66** | -1.7709 | -1.56 | -1.734 | -1.56 | **68** | -1.4771 | -1.529 | -1.441 | -1.625 |
| **69** | -1.6335 | -1.501 | -1.769 | -1.452 | **70** | -2.0414 | -1.125 | -1.187 | -1.891 |
| **72** | -1.3979 | -1.708 | -1.539 | -1.371 | **71** | -1.3617 | -1.6 | -1.753 | -1.543 |
| **73** | -0.8865 | -1.596 | -1.476 | -1.523 | **74** | -3.1139 | -2.754 | -3.029 | -2.58 |
| **75** | -3.2304 | -3.124 | -3.278 | -2.643 | **77** | -2.4914 | -2.609 | -2.565 | -2.687 |
| **76** | -2.0792 | -1.607 | -1.565 | -2.269 | **80** | -2.7782 | -2.483 | -2.47 | -2.414 |
| **78** | -3.301 | -2.68 | -3.074 | -3.228 | **82** | -1.1461 | -1.177 | -1.088 | -1.787 |
| **79** | -2.3617 | -2.66 | -2.694 | -2.231 | **83** | -1.4914 | -1.204 | -1.151 | -1.525 |
| **81** | -1.3979 | -1.133 | -1.195 | -1.291 | **85** | -1.0414 | -1.217 | -1.221 | -1.041 |
| **84** | -0.8388 | -0.886 | -0.632 | -0.678 | **88** | -2.3802 | -2.969 | -3.161 | -2.339 |
| **86** | -1.1139 | -1.159 | -1.089 | -0.78 |  |  |  |  |  |
| **87** | -1.2788 | -1.384 | -1.37 | -1.806 |  |  |  |  |  |

**Figure 3.**
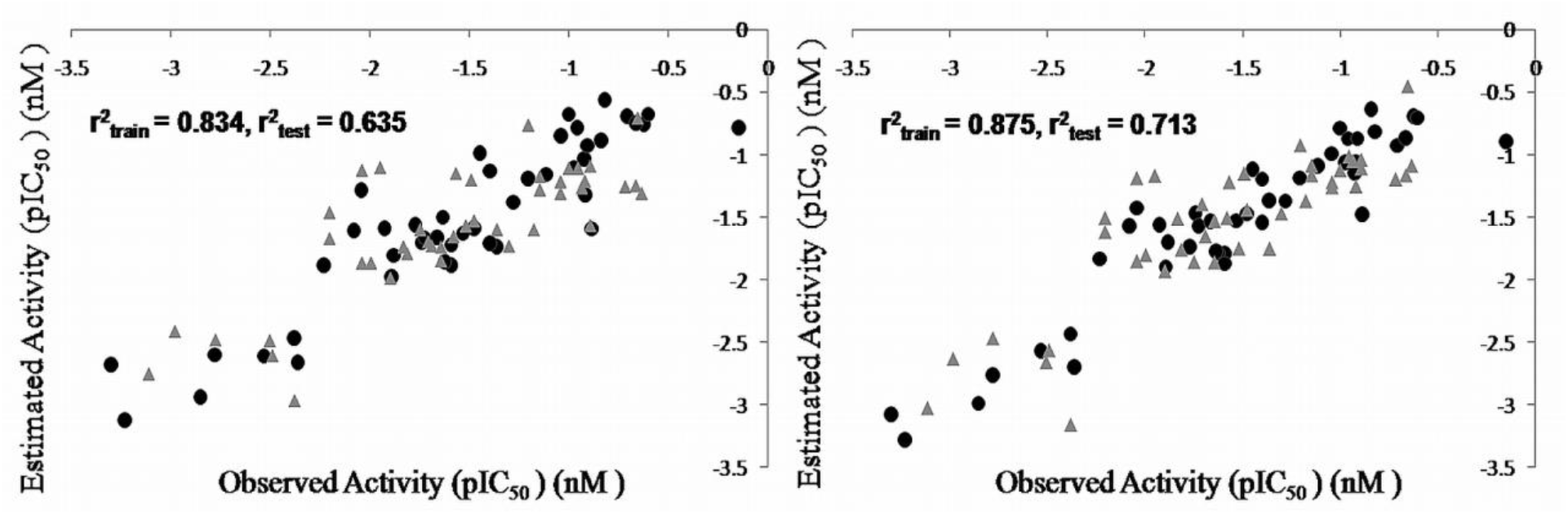
Correlation graph between observed and predicted activities of training set (dots) and test set (triangles) molecules **(A)** CoMFA and **(B)** CoMSIA.

#### 3.1.5 CoMFA and CoMSIA contour maps analysis

The coefficients from CoMFA and CoMSIA models were used to generate 3D contour maps. which determine the vital physicochemical properties responsible for variation in activity and also explore the crucial importance of various substituents in their 3D orientation. The generated Contour maps from the above hypothesis of CoMFA and CoMSIA analyses are shown in Figure. 4A,B,C.

**Figure 4.**
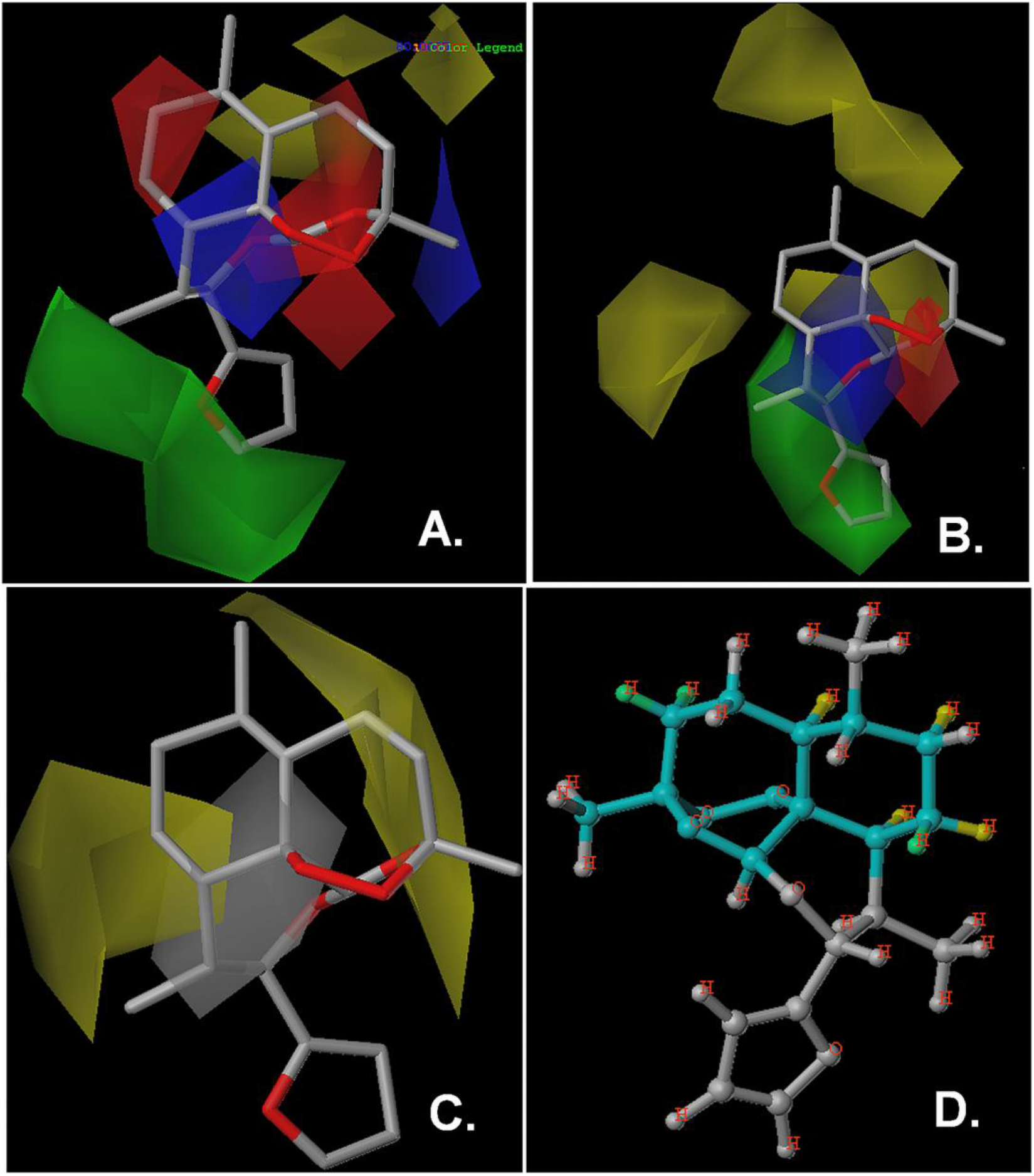
3D-QSAR analysis of substituted 1,2,4-trioxane molecules **(A)** Steric and electrostatic CoMFA contours; **(B)** Steric and electrostatic CoMSIA contours around the most active compound 40. The green and yellow contours indicate sterically favored and disfavored regions respectively while the blue and red contours denote regions that favor electropositive substituent and electronegative substituent respectively; **(C)** PLS hydrophobic contours from CoMSIA analysis around the most active compound 40. The yellow contours indicate hydrophobic favored regions while the white contours denote hydrophobic disfavored regions **(D)** Individual atomic contributions for the activity of the most potent compound **40**. The colors at the red end of the spectrum reflect unfavorable contributions in the order of red > red-orange > orange color, while colors at the green end indicate favorable (positive) contributions in the order of green> green-blue > yellow. Atoms with intermediate contributions are colored in white.

A contrast of the contours obtained from the PLS co-efficient of CoMFA and CoMSIA descriptors yields necessary information about the physicochemical properties of the substituted 1,2,4-trioxane molecules vital in defining the antimalarial activity. An analysis of the CoMFA and CoMSIA contours in terms of common steric and electrostatic parameters around the most active molecule of the dataset **40** signified the importance of sterically favorable region (green color) near the 2-furyl group at C-10 position of the molecule while sterically unfavorable contours (yellow color) have been observed near the region of three carbon chain linked with the C-3 and C-12a position of the trioxane ring, the cyclohexyl ring and the methyl group at C-6 position of the molecule **40** for antimalarial activity (Figure 4A and B). At the same time the CoMSIA based PLS contour analysis of the hydrophobic feature highlighted the importance of the favorable hydrophobic functionalities in the vicinity of the regions adjoining the 3 and 6 position of methyl group as well as regions below the cyclohexyl group of the molecule while the unfourable hydrophobic interactions in the regions near the C-12, 12a, 8a and 9 (Figure 4C). Therefore, it may be inferred that the steric bulk should not be increased at position 3 of the molecule while the hydrophobic groups should be added in the adjacent regions between 3 and 6 positions as well as in the regions below the cyclohexyl group of the molecule while hydrophilicity may be increased in the regions near the C-12, 12a, 8a and 9 of the molecule. Thus this information adds on to the information provided by the DS pharmacophore where though the two aliphatic hydrophobic (HYAl-1 and 2) features were shown but it did not provide any information about the addition or deletion of steric bulk and hydrophobicity in different regions in 3D space of the molecule. Similarly, when the electrostatic contours were considered, both the CoMFA and CoMSIA contours emphasized the importance of the electronegative substituent in the vicinity of the oxygen atoms at 1,2,4 position of the molecule which was corroborated with the two H-bond acceptor features present in the DS based pharmacophore model. However the CoMFA and CoMSIA studies gave the additional information regarding the favorable electropositive functionality in the region adjacent to the 1,2,4-trioxane ring for increasing the antimalarial activity (Figure. 4A and 4B).

### 3.2. HQSAR Study

The role of 1,2, 4 trioxane ring in the artemisinin derivatives for defining the antimalarial activity is well established. Although the CoMFA and CoMSIA studies gave an insight about the quantitative role of chemical features in modulating the antimalarial activity in terms of favorable and unfavorable contours still the HQSAR studies were performed on the same data set to find out the minimal 2D sub-structural requirement for antimalarial activity besides the well known role of 1,2,4 trioxane ring. The HQSAR models were generated using the default fragment size (4–7) combined with various fragment types and various hologram lengths as summarized in Table 2. The model having A, B, C, H, Ch, and DA fragments with r^2^_ncv_ value of 0.873 at 6 components and 199 hologram lengths was selected. In order to further analyse it, the data set was divided into the same training and test set as in CoMFA and CoMSIA studies to access the predictive values of the model. The observed r^2^_ncv_ value 0.848 and 0.910 between experimental and predicted pIC_50_ of the training and test set respectively further signified the quality of the model. The predictive pIC_50_ values of the training as well as test set molecules based on the HQSAR model are listed in the Table 4 and plotted in Figure 5.

**Table 4.** Results of HQSAR analyses for various fragment distinctions on the key statistical parameters using fragment-size default (4–7).

| Model | Fragment<sup>a</sup> | $r^2_{cv}$ | SEP | $r^2_{ncv}$ | HL | N |
| --- | --- | --- | --- | --- | --- | --- |
| 1 | A/B/C | 0.585 | 0.369 | 0.716 | 257 | 4 |
| 2 | A/B/C/H | 0.573 | 0.392 | 0.675 | 151 | 3 |
| 3 | A/B/C/H/Ch | 0.557 | 0.359 | 0.713 | 151 | 4 |
| 4 | A/C/H/Ch/DA | 0.578 | 0.267 | 0.855 | 199 | 6 |
| 5 | **A/B/C/H/Ch/DA** | **0.572** | **0.249** | **0.873** | **199** | **6** |
| 6 | A/B/Ch/DA | 0.571 | 0.378 | 0.697 | 97 | 3 |
| 7 | A/B/C/Ch/DA | 0.500 | 0.401 | 0.660 | 151 | 3 |
| 8 | A/C/Ch/DA | 0.483 | 0.451 | 0.566 | 307 | 2 |

SEP: standard error of prediction, HL: hologram length, N: optimal number of components, <sup>a</sup>Fragment distinction; A: atom, B: bond, C: connection, H: hydrogen, Ch: chirality, DA:donor and acceptor.

**Figure 5.**
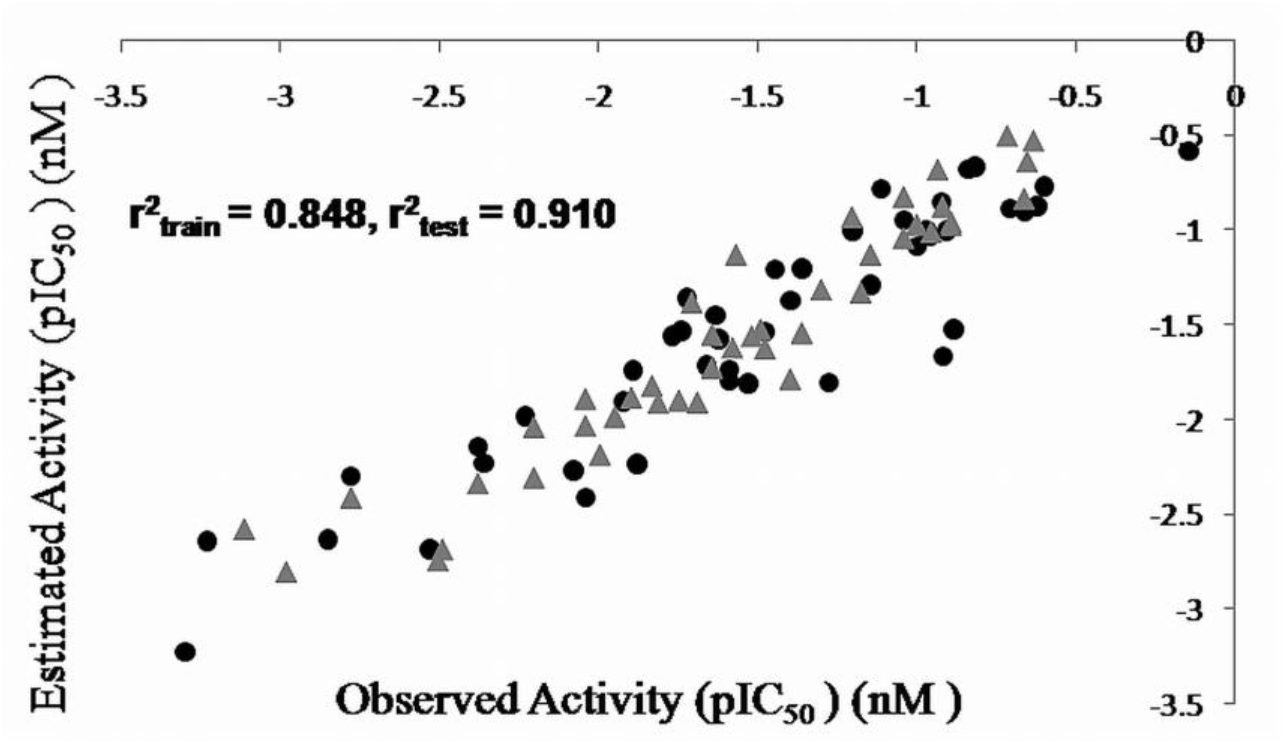
Correlation graph between observed and predicted activities of training set (dots) and test set (triangles) molecules based on HQSAR model.

An attractive property of HQSAR technique is that it provides straightforward clues about the individual atomic contributions to the biological activities through the use of different color codes. The color coding is based on the activity contribution of the individual atoms of the molecules. The individual atomic contribution of the most potent compound **40** is shown in Figure 4D where the cyclohexy group and ring formed by the 3C chain linkers both attached with trioxane ring define the antimalarial activity besides the well known 1,2,4 trioxane ring. These findings also corroborates with the CoMFA, CoMSIA and DS based pharmacophore studies where these chemical group were found favorable in defining the antimalarial activity.

## 4. Conclusion

The present study describes a successful application of combination of three different computational approaches to identify essential structural requirements in 3D chemical space for the modulation of the antimalarial activity of substituted 1,2,4-trioxanes. Each approach has its own advantages and disadvantages. The CoMFA and CoMSIA have been applied successfully to rationalize the 3D space in diverse substituted 1,2,4-trioxanes in terms of their steric, electrostatic, and hydrophobic interaction for their antimalarial activity. The developed models showed good statistical significance in internal validation (q^2^, group cross-validation and bootstrapping) and performed very well in predicting antimalarial activity of 43 substituted 1,2,4-trioxanes in the test set. The CoMFA, CoMSIA models not only provided the information about the favorable regions as reported in DS based pharmacophore but also give the information about the unfavorable regions in defining the potency. In addition the superiority of QuantitativePBA over GMCBA has been established. Further, the HQSAR studies based on the same training set also provided the 2D sub-structural requirements and showed good statistical significance in internal validation (r^2^_cv_, r^2^_ncv_) as well as predicted very well the antimalarial activity of the test set compounds. Thus, the three layered QSAR studies *viz*. Pharmacophore, CoMFA, CoMSIA and HQSAR may be useful for designing new substituted 1,2, 4-trioxane with potent antimalarial activity.

## Acknowledgements

The authors are thankful for financial assistance in the form of fellowship by Indian Council of Medical Research (ICMR) (A.K.G.), and to Mr. A. S. Kushwaha for technical assistance. CDRI communication No. allotted to this manuscript is AKS12.

